# The monomeric conformational ensembles of A*β*40 and A*β*42 encode their differential amyloid aggregation propensity

**DOI:** 10.64898/2026.01.30.702803

**Authors:** Irene Cadenelli, Andrea Ciccolo, Andrea Tagliabue, Giulia Rossi, Valeria Conti-Nibali, Davide Bochicchio

## Abstract

A*β*40 and A*β*42 peptides differ by just two C-terminal residues, yet they display strikingly different aggregation and toxicity profiles. Whether this distinction is already encoded at the monomer level is still under debate. Here, we combine extensive all-atom simulations in explicit solvent, well-tempered metadynamics, and a tailored consensus cluster analysis to compare the monomeric ensembles of the two isoforms under identical conditions. Both peptides populate broad, coil-like conformational distributions; however, A*β*42 shows systematically higher *β*-structure propensity, especially in the C-terminal region, and samples more extended conformations with higher hydrophobic exposure compared to A*β*40. These results support a mechanistic link between sequence-encoded monomer conformational preferences and the differential amyloidogenicity of the two isoforms, highlighting monomer-level determinants of A*β*42’s distinct aggregation behavior.

## Introduction

Alzheimer’s Disease (AD) is a widespread neurodegenerative disorder characterized by a progressive decline of cognitive function, dysfunction, and neuronal loss [1, 2, 3, 4], and currently ranks among the leading causes of death and disability worldwide, particularly in older populations [5]. The main hallmarks of AD are the extracellular deposits of amyloid fibrils formed by amyloid-*β* (A*β*) peptides, predominantly by the A*β*40 and A*β*42 isoforms. Although these two isoforms differ by only two C-terminal residues, they display markedly distinct aggregation kinetics, morphologies, and toxicities [6, 7]. Compared with A*β*40, A*β*42 is more abundant in mature amyloid fibrils, even though A*β*40 is produced in higher amounts from the amyloid-beta precursor protein [8]. In addition, A*β*42 aggregates more rapidly and is more capable of damaging neuronal membranes by aggregating into pore-forming oligomers [6, 7].

Amyloid oligomers (4–10 monomers) can be significantly more toxic than micrometer-long, mature fibrils. Fibril-catalyzed *secondary nucleation*—the formation of new nuclei on the surface of existing fibrils—can dominate aggregate proliferation and enhance the generation of oligomeric toxic species, especially for A*β*42 [6, 7, 9, 10].

At the monomer level, both A*β*40 and A*β*42 are intrinsically disordered peptides (IDPs) that populate very broad conformational ensembles in aqueous solution [11]. Single-molecule spectroscopy and all-atom simulations indicate highly dynamic, coil-like distributions with rapid interconversion among conformers [12]. The vast heterogeneity of possible structures at the monomer level is reflected in a certain degree of polymorphism of amyloid fibrils. However, the number of structures identified for amyloid fibrils is substantially lower than the number of monomeric conformers. Therefore, whether and how subtly sequence-encoded monomer features can bias downstream aggregation pathways into different structures remains a key open problem. A recent work demonstrated that, despite amyloid polymorphism, the sequence regions most often forming the structural core of amyloid fibrils are those exhibiting high intrinsic *β*-sheet propensity in the monomer [13]. This observation suggests that, in principle, the aggregation propensity of different isoforms could be anticipated from their distinct monomeric conformational preferences in solution. Experimentally, however, isolating and characterizing truly monomeric A*β* species remains extremely challenging, as IDPs rapidly self-associate even at sub-micromolar concentrations, populate fast monomer–oligomer equilibria, and quickly adsorb onto experimental substrates [14].

In this context, molecular simulations with atomistic resolution have become essential tools for understanding these complex systems. Methodological developments now enable a more reliable characterization of the disordered states of proteins. In particular, protein force fields refined for IDPs (e.g., CHARMM36m, a99SB-disp) have substantially improved agreement with SAXS, NMR, and smFRET benchmarks for unfolded and disordered proteins [15, 16, 17]. In parallel, enhanced-sampling strategies, such as well-tempered metadynamics, combined with time-independent reweighting, allow reconstruction of unbiased thermodynamics over slow collective variables (CVs), providing statistically robust estimates of conformational populations and free-energy basins for flexible peptides [18, 19].

Furthermore, machine-learning approaches have begun to enhance our understanding of the conformational landscapes and transition pathways of amyloid beta peptides in solution. Notable examples include neural-network Markov models, such as VAMPNet/CoVAMPnet [20, 21], which learn soft state decompositions and kinetics, as well as graph-based variants that scale and regularize these analyses on large datasets [22], and deep generative models that sample IDP-consistent conformers [23, 24, 25].

Prior simulation work on A*β* peptides showed clearly that, under physiological conditions, A*β*40 and A*β*42 populate an ensemble of random-coil–like conformations, even if secondary structures along the sequence are transiently sampled [26, 27]. Bias-Exchange Molecular Dynamics simulations guided by NMR data have revealed an inverted free-energy landscape: the global minimum corresponds to an ensemble of highly disordered conformations, whereas regions of higher free energy are associated with more structured states displaying an increasing content of secondary structure elements [28]. For both isoforms, the *α*-helix content is generally lower than the *β*-sheet content, consistent with circular dichroism data indicating an *α*-helix content of ≈9% and a *β*-content of 12-25% [29, 30]. However, it has also been suggested that there are isoform-specific differences, especially in the balance between compact and extended conformations, as well as in the organization of hydrophobic contacts and C-terminal *β*-propensities [31]. Still, consensus has been limited by well-known force-field and water-model biases affecting the dimensions and secondary-structure propensities of intrinsically disordered proteins [17, 16, 32], as well as by the lack of consistency in force fields, simulation protocols, and analysis pipelines across computational studies of A*β*40 and A*β*42 in solution [33, 34, 35]. In this study, we adopt a systematic computational strategy to determine whether structural differences between A*β*42 and A*β*40 already emerge at the monomer level. Using extensive allatom simulations in explicit solvent with an IDP-calibrated force field, combined with advanced enhanced sampling, rigorous reweighting, and a clustering framework that robustly identifies dominant conformational families across statistically equivalent trajectory subsets, we compare the monomeric ensembles of the two isoforms under identical conditions. This approach allows us to resolve isoform-specific differences in secondary structure propensities, hydrophobic exposure, and contact patterns that are consistent with known fibril-forming interfaces [36, 37, 13]. Our results suggest that sequence-encoded monomeric conformational preferences contribute to the distinct aggregation behavior of A*β*40 and A*β*42.

## Results

We characterized and compared the monomeric conformational ensembles of A*β*40 and A*β*42 using all-atom simulations with the CHARMM36m force field [15] in explicit water under near-physiological conditions (0.15 M NaCl, 310 K). As a first step, we reconstructed the free energy surfaces (FESs) of the two isoforms. We then characterized the resulting conformational ensembles in terms of residue-resolved secondary-structure propensities, global compaction, and hydrophobic exposure, as well as the dominant metastable families identified through a tailored clustering procedure. See section *Methods* and the Supporting Information (SI) for simulation details.

### Global free energy surfaces

To obtain a global picture of the conformational preferences of the two monomers in water, we reconstructed the free-energy surfaces (FESs) of A*β*40 and A*β*42 using well-tempered metadynamics. We used two collective variables (CVs) describing secondary structure content: the *α*- and *β*-content, implemented in PLUMED as `ALPHARMSD` and `ANTIBETARMSD` [38] and described in detail in the *Methods* section. Convergence is a well-known critical aspect in the study of IDPs, whose conformational landscapes are broad and only weakly funneled. We monitored the stability of the sampling achieved in these simulations by monitoring the time evolution of the free energy difference between relevant FES regions (see SI for more details). The results indicate that the sampling is sufficiently stable to support the comparative analysis presented below for A*β*40 and A*β*42.

In order to present the results in a more easily interpretable collective variable space, the original CVs values were rescaled to yield approximate residue counts for *α*-helical (*n*_*α*_) and *β*-sheet (*n*_*β*_) structures. The FESs were therefore represented in the (*n*_*α*_, *n*_*β*_) plane for each isoform. The resulting two-dimensional FESs and their corresponding one-dimensional projections are shown in Figure 1.

**Figure 1:**
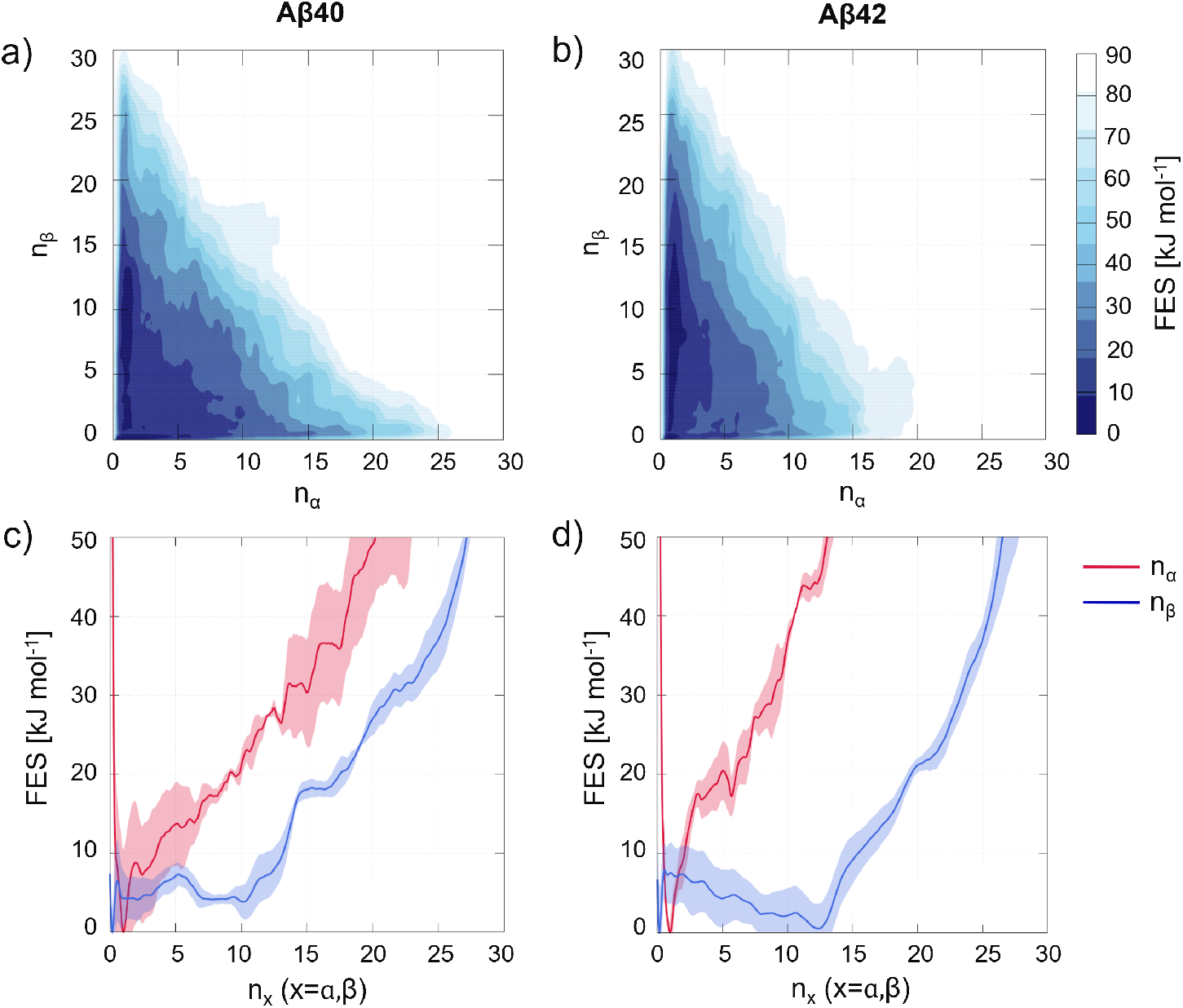
A*β*40 and A*β*42 free energy surfaces. FES for (a) A*β*40 and (b) A*β*42 in the (*n*_*α*_, *n*_*β*_) plane. The variables *n*_*α*_ and *n*_*β*_ approximate the number of residues in *α*-helical and *β*-like conformations, respectively, and highlight the different balance between helical and *β*-rich states in the two isoforms. Below, one-dimensional free energy profiles for (c) A*β*40 and (d) A*β*42, projected onto *n*_*α*_ and *n*_*β*_. Solid lines show the reweighted free energy profiles, with shaded regions indicating the standard error of the mean across the three independent replicas.

The FESs of both peptides reflect their intrinsically disordered nature, displaying the characteristic inverted funnel shape in which highly disordered states occupy the lowest free-energy region, while more ordered conformations enriched in secondary structure elements correspond to higher free energy values [28, 39]. Both isoforms span broad, multimodal conformational spaces. The most populated regions correspond to poorly structured, coil-like states, with additional basins associated with significant *β*-sheet content.

Across both peptides, *β*-like conformations are generally favored over *α*-helical ones. A*β*42 samples *β*-rich regions more extensively than A*β*40, whereas A*β*40 explores the landscape more broadly, visiting low and medium *α*-content regions more frequently, although these remain high at free energy values.

### Local secondary structure patterns and sequence-dependent differences

Per-residue secondary structure probabilities (Figure 2) were obtained by reweighting (see *Methods*) and analyzing each trajectory with DSSP (version 3.1.4) [40]. For every frame and every residue, DSSP assigns one among eight local conformations–i.e., 3_10_ helix, *α*-helix, *π*-helix, polyproline II helix, extended *β*-strand, isolated *β*-bridge, turn, and bend–with remaining states reported as coil. Each assignment was encoded as a binary indicator and reweighted frame by frame, yielding unbiased probabilities *p*_*i*_(*s*) for all DSSP states *s* at each residue *i*. In Figure 2, the probability distributions {*p*_*i*_(*s*)} across the eight DSSP categories are shown for each residue.

**Figure 2:**
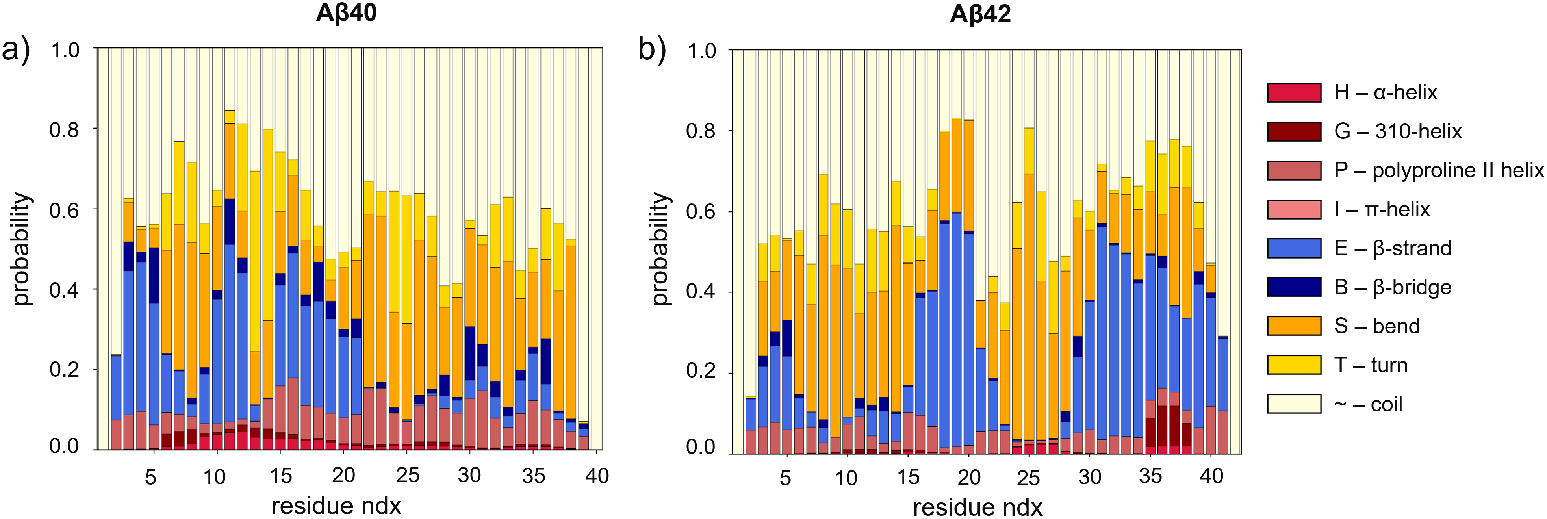
Per-residue secondary structure probabilities for A*β*40 (a) and A*β*42 (b). Each vertical bar corresponds to a residue, and the colored stacked segments represent the full probability distribution {*p*_*i*_(*s*)} across eight DSSP categories. The total height of each bar is one, providing a direct readout of the relative sampling of each secondary structure type in the unbiased ensemble.

For both peptides, as expected, coil-like states dominate, but pronounced sequence-dependent differences emerge in the pattern and localization of *β*-strand formation.

A*β*40 displays three well-separated *β* peaks in the first half of the sequence (approximately residues 2–7, 9–13, and 15–21), each sharp and localized, with intervening residues reverting almost entirely to coil or turn states. Beyond residue ∼21, in the C-terminal region, *β* structures are only sparsely observed.

A*β*42, by contrast, shows a distinctly different organization. In the N-terminal half, the first *β* peak (residues 2–7) is present but weaker than in A*β*40, and the second peak (9–13) is nearly absent. The third peak (15–21) is instead present and relevant. The most striking feature is a broad, continuous, and strongly stabilized *β*-rich region extending approximately from residue 29 to the C-terminus. This persistent C-terminal *β*-segment has no counterpart in A*β*40 and is consistent with independent simulations and NMR-refined ensembles highlighting enhanced *β*-structure in the 30–36 and 39–41 regions of A*β*42. [41, 42, 43].

Additional DSSP features contextualize these differences. Turns and bends are abundant in both isoforms and preferentially cluster near the boundaries of the *β* regions. Helical conformations are overall rare but sampled slightly more frequently in A*β*40, indicating a more heterogeneous and structurally flexible ensemble.

We conducted a complementary analysis by calculating intra-peptide contact maps (Figure 3) by reweighting and analyzing each trajectory with the MDAnalysis Python library [44]. Pairwise residue distances were calculated considering only C_*α*_ atoms, and a cutoff of 0.8 nm was applied to define residue–residue contacts. The contact maps are consistent with the trends observed in the secondary-structure analysis (Figure 2) and offer a detailed, quantitative characterization of the peptides’ dynamic organization.

**Figure 3:**
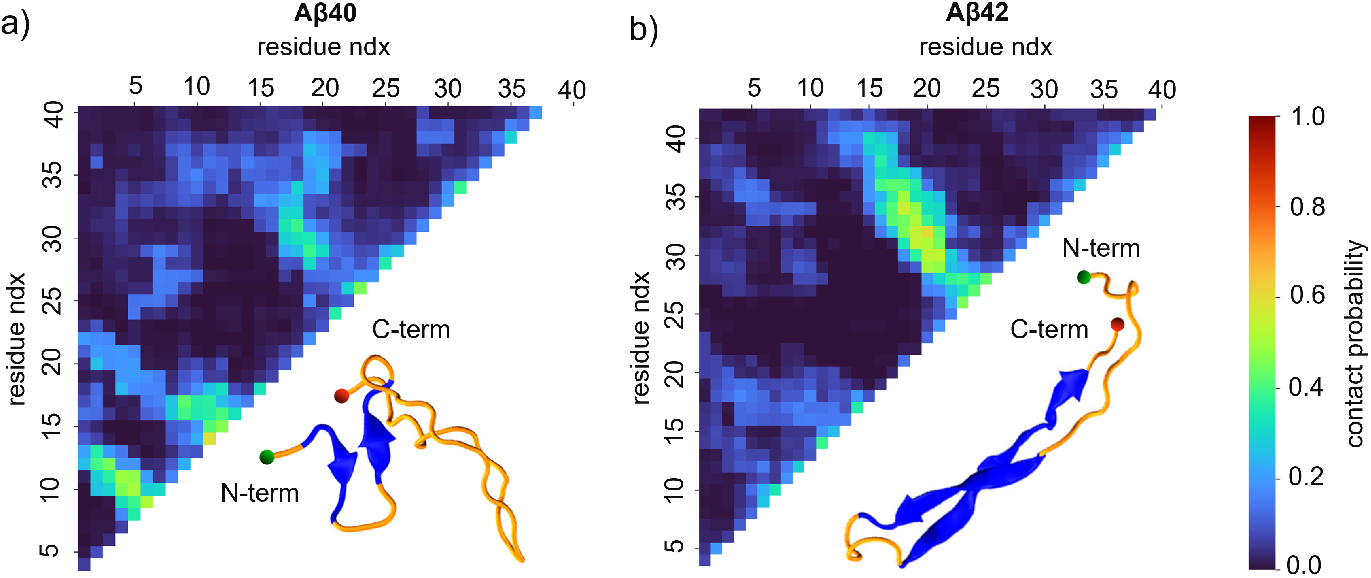
Intra-peptide contact maps and representative structures sampled for (a) A*β*40 and (b) A*β*42 (b). Each square represents the contact probability between pairs of residues. The main diagonal is masked to exclude trivial contacts between sequentially adjacent residues. Beside the contact maps, an example representative snapshot of the corresponding peptide is shown. The N-terminal and C-terminal ends of both peptides are indicated by green and red beads, respectively. Blue arrows highlight antiparallel *β*-sheets.

For A*β*40 (Figure 3a) the highest contact probabilities are localized within the first half of the peptide sequence (residues 2–16), consistent with the local folding and stabilization of short and transient compact structures. There is another shallow contact region involving residues in the central part of the peptide; in contrast, the C-terminal region (29–40) shows no significant propensity to establish persistent contacts with other segments of the peptide.

Conversely, for A*β*42 (Figure 3b), the C-terminal region exhibits the highest contact probabilities, particularly through interactions with the central hydrophobic core (residues 17–21). These observations align with previous studies [45, 46] that relate the enhanced *β*-sheet propensity, the presence of two additional residues, and the higher structural rigidity of the A*β*42 C-terminal, and suggest a higher aggregation propensity compared to the more flexible A*β*40.

### Compaction and solvent exposure

A*β*40 and A*β*42 exhibit similar *R*_*g*_ distributions, both dominated by a main peak in the 1.0– 1.5 nm range (Figure 4a,d), in line with previous all-atom simulations of A*β* monomers in explicit solvent [34, 47]. This main peak corresponds to compact conformations representing the dominant population of both isoforms. However, A*β*42 displays a more evident tail extending up to ∼2.5–3.0 nm, whereas A*β*40 decays more rapidly beyond 2 nm.

**Figure 4:**
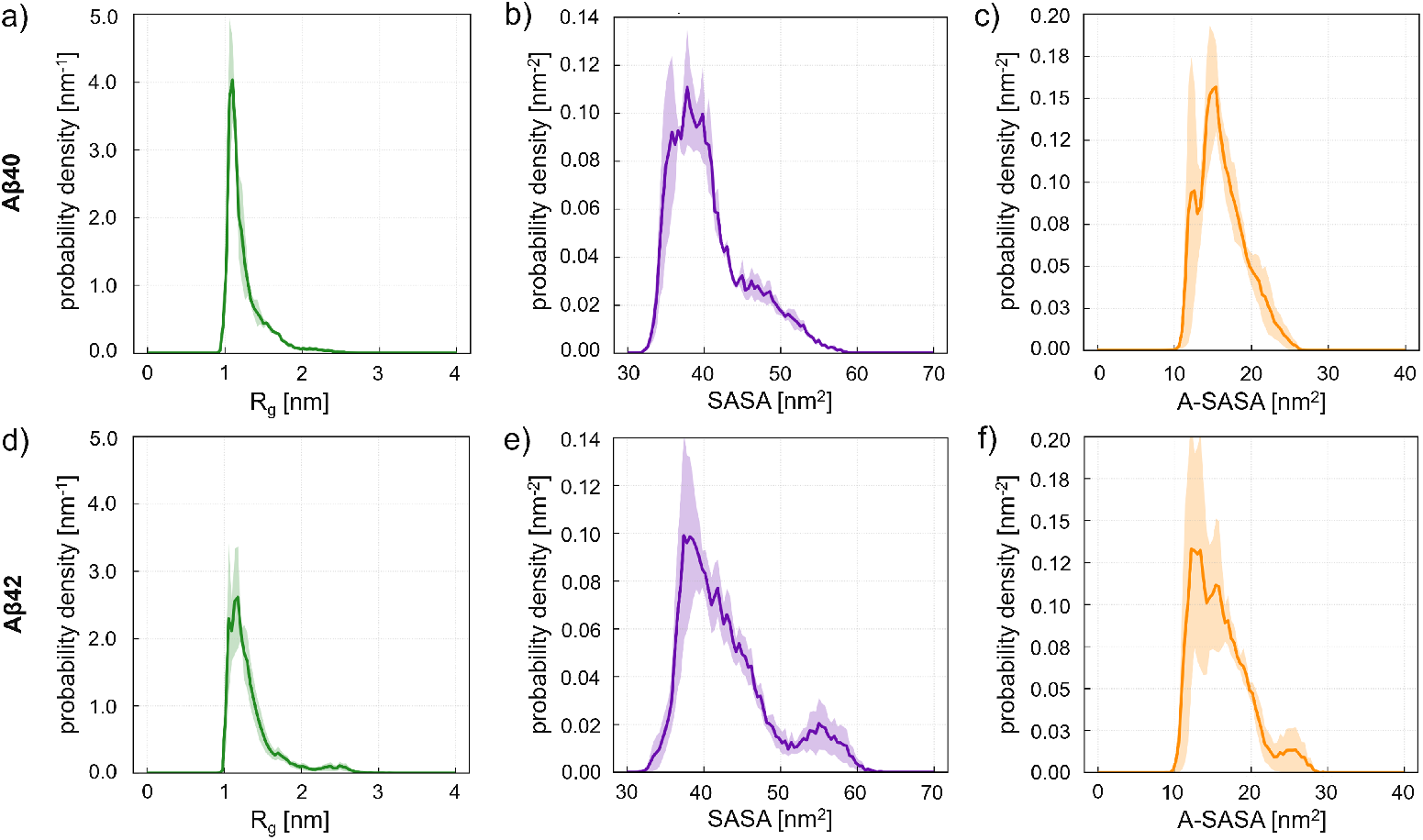
Conformations and exposure to solvent of (a–c) A*β*40 and (d–f)A*β*42. (a,d) Probability distributions of the radius of gyration (*R*_*g*_). (b,e) Total solvent-accessible surface area (SASA). (c,f) Apolar SASA (A-SASA). Shaded regions represent the standard error of the mean across independent replicas.

Consistent with these observations, the SASA distributions of both peptides show a primary peak around ≈40 nm^2^ (Figure 4b,e). For A*β*40, this peak is followed by a gentle, featureless shoulder at larger SASA values, reflecting a continuum of moderately more exposed conformations. In contrast, A*β*42 exhibits a second, distinct peak in the 50–60 nm^2^ range, marking a well-defined population of highly solvent-exposed structures.

In addition to the total SASA, apolar SASA (A-SASA) was computed from the exposure of hydrophobic residues according to the Roseman scale [48] (See *Methods* for further details). A similar trend emerges for A-SASA (Figure 4c,f). Both peptides show a main peak between 15 and 20 nm^2^, corresponding to compact states with partially buried hydrophobic residues. Only A*β*42 displays an additional shoulder or minor peak between 20 and 30 nm^2^, indicating access to conformations with substantially larger hydrophobic exposure.

### Conformational clustering

To resolve the dominant conformational families underlying the free energy surfaces of A*β*40 and A*β*42, we developed an *ad hoc* clustering framework. Standard clustering approaches often struggle when applied to intrinsically disordered peptides, whose free energy landscapes are characterized by broad and partially overlapping basins. When projected onto a low-dimensional space defined by a small set of CVs, these basins often appear as clusters of configurations with highly heterogeneous local densities. The clustering strategy adopted here was therefore specifically designed to address these challenges by combining density-based detection across multiple scales with a consensus-based reconciliation of independently clustered trajectory subsets. A schematic overview of the workflow is shown in Figure 5, while the full workflow is explained in the following.

**Figure 5:**
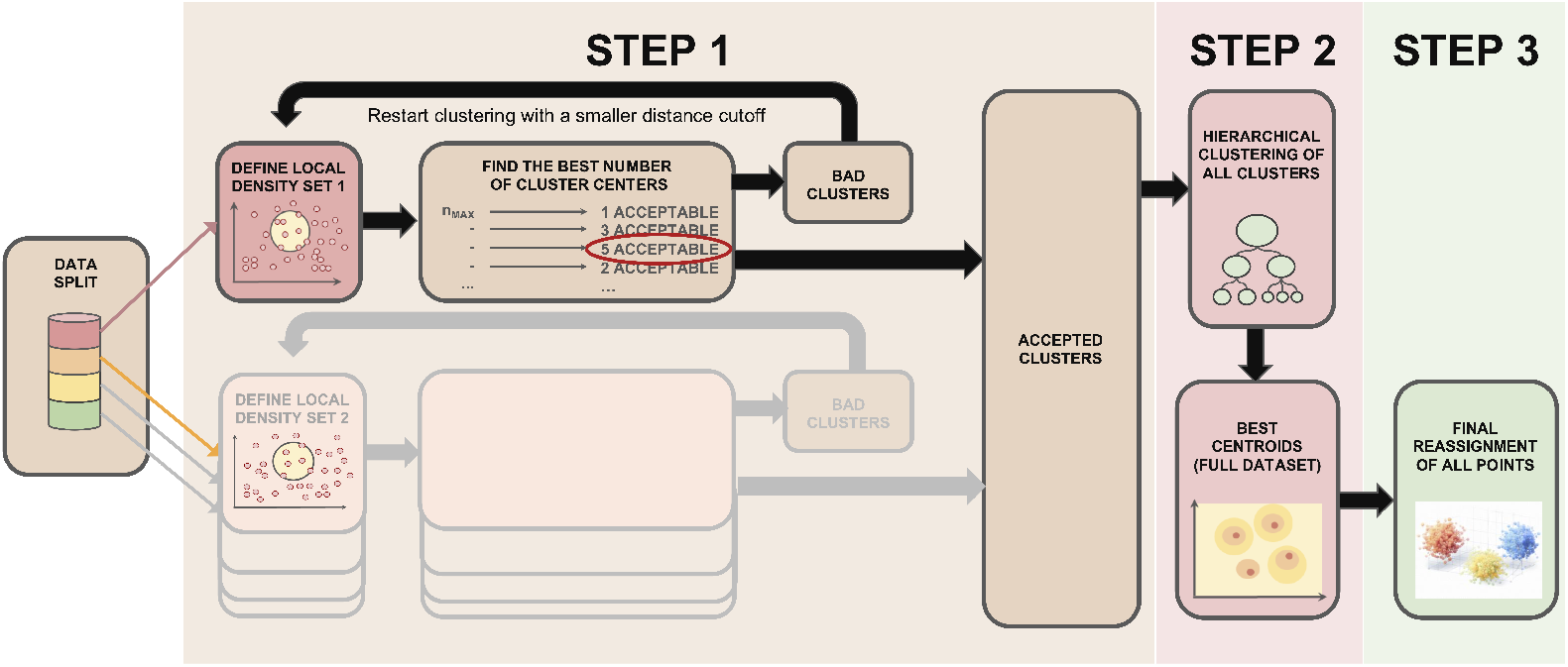
Schematic overview of the conformational clustering workflow. The procedure, described in detail in the main text, combines iterative density-based clustering and consensus consolidation to identify robust conformational families in heterogeneous ensembles.

#### Consensus clustering workflow

Clustering was performed in the space of four structural descriptors displaying a strong multimodality in the reweighted ensemble: *n*_*α*_, *n*_*β*_, the radius of gyration *R*_*g*_, and total SASA. This choice ensured sensitivity to major conformational basins. Only once the final clusters were obtained, to further characterize them, we added four additional descriptors to the analysis, namely the apolar solvent-accessible surface area (A-SASA), the polar solvent-accessible surface area (P-SASA), the number of intra-peptide hydrogen bonds (*n*_HB− IP_), and the number of peptide–solvent hydrogen bonds (*n*_HB−PS_). Furthermore, to improve statistical robustness and control memory usage, trajectories were merged and partitioned into random subsets of about 3 ×10^4^ frames. We verified, by comparing them, that the distributions of structural descriptors were essentially indistinguishable across the different subsets.

The first step of our clustering approach (STEP 1 in Figure 5) applies a parallel workflow to each subset. The workflow is based on the standard DPC algorithm [49], which identifies cluster centers as points of simultaneously high density and large distance from any denser point.

DPC is applied recursively, each time reducing the number of requested clusters from *n*_*max*_ to 2. As a result of this scan, the optimal *n* is identified as that generating the highest number of acceptable clusters (silhouette score ≥*s*_*min*_ and size *N* ≥*N*_*min*_ frames). Acceptable clusters are then removed from the subset sample, and saved. The discarded frames are re-submitted to DPC with a reduced distance cutoff (see Methods for details). The procedure is iterated until no acceptable clusters are identified. At the end of STEP 1, it is possible that some frames remain unassigned.

In STEP 2 (Figure 5), all acceptable clusters identified at STEP 1 from all the trajectory subsets are subject to a hierarchical clustering procedure, based on the Wasserstein (Earth Mover’s) distance and agglomerative hierarchical clustering with average linkage [50], as better detailed in the *Methods* section. The output of STEP 2 is the final list of centroids.

In STEP 3, all the original trajectory frames, including those that were not assigned in STEP 1, are re-assigned to their closest centroid.

The relevance of each conformational family was computed *a posteriori* by incorporating the reweighting factors, thus providing an unbiased estimate of its equilibrium population. The specific parameters used in the clustering pipeline, together with additional technical details, are provided in the *Methods* section, while the SI reports the full structural characterization for all clusters.

#### Outcomes

Figures 6 and 7 summarize the structural characterization of the dominant conformational families identified by the consensus clustering for A*β*40 and A*β*42. Figure 6 compares clusters using the descriptors defining the clustering space, while Figure 7 provides an additional *a posteriori* characterization based on complementary observables–i.e., A-SASA, P-SASA,*n*_HB_ −_IP_, and *n*_HB_ −_PS_. In both figures, clusters are reported in decreasing order of relevance. Throughout the manuscript, cluster indices (e.g., Cluster 1, Cluster 2) refer to the order of relevance for the corresponding peptide, as defined in Figs. 6 and 7.

**Figure 6:**
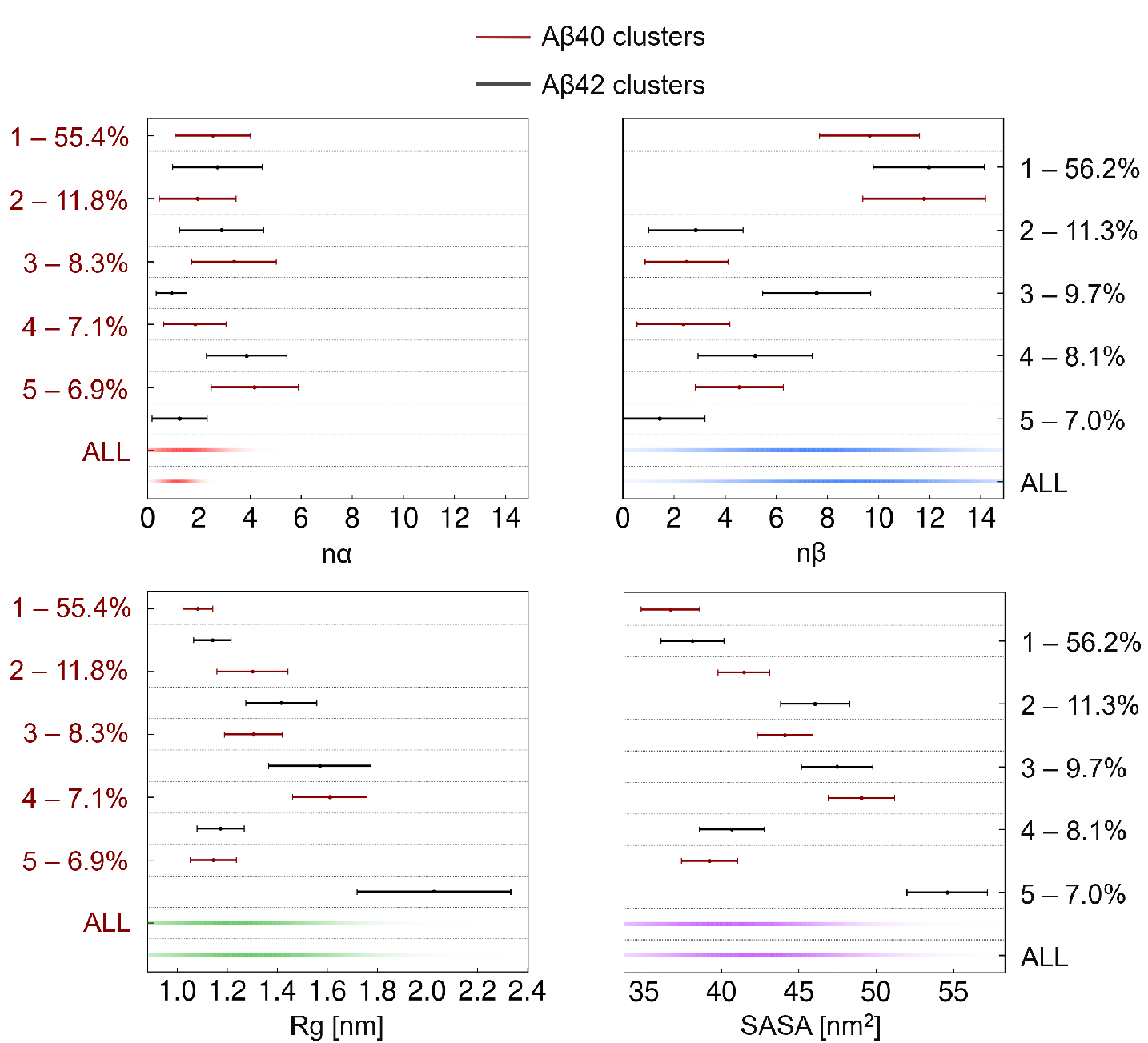
Characterization of the dominant structural families in terms of the structural descriptors used in the consensus clustering analysis. Panels (a–d) report, respectively, the structural descriptors *n*_*α*_, *n*_*β*_, *R*_*g*_, SASA, which define the clustering space. In each panel, the five most relevant clusters of A*β*40 and A*β*42 are shown, ordered by decreasing relevance, with the corresponding reweighted population reported in parentheses. For each cluster and descriptor, the weighted mean value is reported together with a horizontal bar representing the width of the reweighted distribution. For reference, background distributions corresponding to the full reweighted ensembles of A*β*40 and A*β*42 are shown, with color intensity proportional to the probability density.

**Figure 7:**
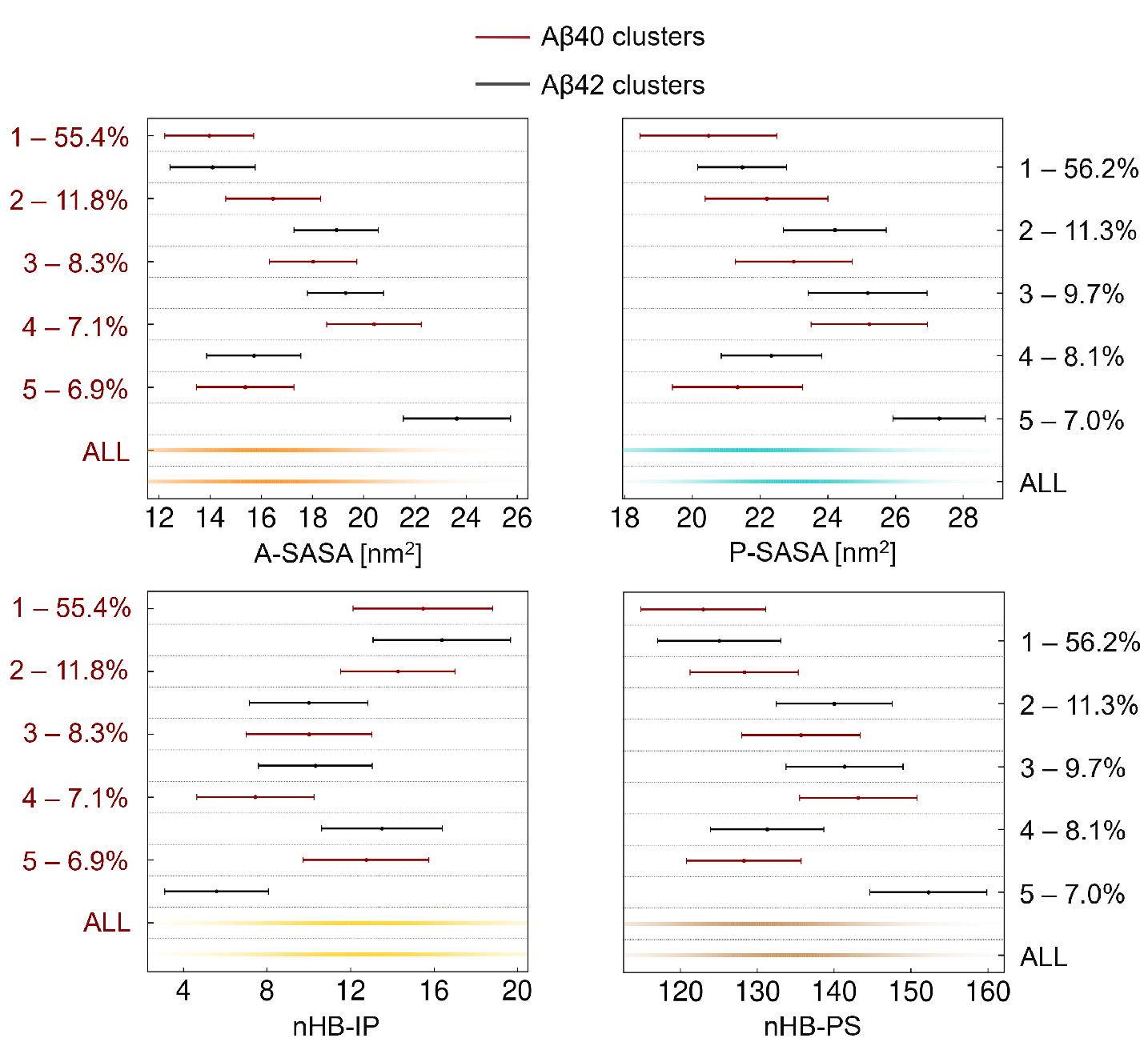
Characterization of the dominant structural families in terms of structural descriptors *excluded* from the consensus clustering analysis. Panels (a–d) report, respectively, A-SASA, P-SASA, *n*_HB− IP_, and *n*_HB−PS_, computed *a posteriori* and not used for clustering. All other conventions are as in Figure 6.

The dominant conformational families identified by the consensus clustering reveal clear structural correspondences as well as systematic differences between A*β*40 and A*β*42. A detailed comparison of the most relevant clusters highlights how the two isoforms populate similar basins with distinct degrees of compactness, secondary structure content, and solvent exposure:

#### Dominant compact *β*-rich basin

Cluster 1 in both peptides corresponds to compact, *β*-rich conformations that dominate the equilibrium ensemble. Notably, A*β*42 exhibits a slightly higher degree of internal organization than A*β*40, as reflected by its larger number of intra-peptide hydrogen bonds.

#### Extended *β*-rich conformations

A clear correspondence emerges between Cluster 2 of A*β*40 and Cluster 3 of A*β*42. These states represent more extended and slightly less *β*-rich conformations with respect to Cluster 1. In this regime, A*β*40 displays a higher *β*-structure content, whereas A*β*42 adopts more expanded conformations, characterized by increased *R*_*g*_, SASA, A-SASA, and P-SASA. Consistently, A*β*40 retains a larger number of intra-peptide hydrogen bonds, while A*β*42 interacts more extensively with the solvent.

#### Highly disordered compact states

A second pair of corresponding states is captured by Cluster 3 of A*β*40 and Cluster 2 of A*β*42. Both clusters are highly disordered while remaining relatively compact; nevertheless, A*β*42 adopts more open conformations and displays higher SASA values.

#### Mixed and weakly *α*-structured compact ensembles

Cluster 5 of A*β*40 and Cluster 4 of A*β*42 represent the only basins with detectable, albeit low, *α*-helical content. These clusters exhibit comparable *β*-levels and correspond to mixed, compact, and internally organized conformational ensembles.

#### Expanded and solvent-exposed disordered states

Finally, Cluster 4 of A*β*40 and Cluster 5 of A*β*42 correspond to more disordered and expanded states with increased solvent exposure. The A*β*42 basin is significantly more extended and solvent-exposed, in agreement with the bimodal behavior observed in the SASA and A-SASA distributions (Figure 4).

## Discussion and Conclusion

By combining molecular dynamics simulations with enhanced sampling techniques, this work aimed to determine whether monomer-level conformational biases could already provide a structural basis for the distinct aggregation propensities observed for A*β*40 and A*β*42. While many aspects of the conformational ensembles of these two peptides have been examined previously, here we provide a direct comparison obtained under identical simulation and analysis conditions (same sampling strategy, force field, and post-processing pipeline).

The reconstructed free energy surfaces (Figure 1) show that both peptides predominantly populate disordered, coil-like ensembles, in agreement with previous computational studies [28, 39]. Nevertheless, clear differences emerge in their secondary structure balance. A*β*42 exhibits a higher probability of adopting *β*-like conformations relative to *α*-helical ones, exceeding the corresponding tendency observed in A*β*40. Conversely, A*β*40 samples a broader range of conformations, including more frequent excursions into transient *α*-helical regions. These global trends indicate a subtle but systematic shift in the balance between disorder and transient structural order that is already encoded at the monomeric level.

The residue-resolved DSSP analysis (Figure 2 refines this picture by identifying the sequence regions responsible for these global differences. In particular, the extended C-terminal *β* domain observed in A*β*42 closely mirrors experimentally established distinctions between the fibril cores of the two peptides. Solid-state NMR and cryo-EM studies consistently identify residues 30–42, together with a central segment around residues 10–20, as major contributors to the cross-*β* core in A*β*42 fibrils, whereas A*β*40 incorporates C-terminal residues only weakly, or not at all, under comparable conditions [36, 51, 37, 52]. Recent meta-analyses by Errico *et al*. [13] further show that, across dozens of *in vitro* and brain-derived A*β* fibril polymorphs, dominant cross-*β* cores almost invariably involve overlapping C-terminal segments, despite their high structural diversity. Although the present simulations focus on monomeric species, the presence of such motifs suggests that the A*β*42 monomeric ensemble samples conformations that are structurally consistent with aggregation-prone features identified experimentally. In contrast, the more fragmented and heterogeneous *β*-strand distribution observed in A*β*40 is associated with a conformational landscape that appears less prone to forming the contiguous *β* segments characteristic of mature fibrils. Consistently, the intra-peptide contact maps reveal an isoform-specific reorganization of tertiary contacts (Figure 3): A*β*42 displays enhanced contact probability between the C-terminal segment and the central hydrophobic core, whereas A*β*40 shows contacts mainly confined to the N-terminal/central region with little persistent involvement of its C-terminus.

Global shape and solvent exposure provide a complementary perspective on the monomeric ensembles (Figure 4). In our simulations, A*β*42 displays a slightly broader distribution of the radius of gyration compared to A*β*40, together with a distinct secondary peak in the SASA and A-SASA distributions associated with extended, highly solvent-exposed conformations. By contrast, the corresponding distributions for A*β*40 decay more rapidly toward larger radii and higher solvent exposure. These A*β*42-specific shoulders indicate that, while both isoforms share similarly most-probable compact states, A*β*42 more readily samples conformations with larger surface area and greater hydrophobic exposure.

Previous experimental and computational studies have reported partially divergent views on the conformational differences between A*β*40 and A*β*42. Single-molecule FRET measurements combined with molecular dynamics simulations by Meng *et al*. [12] suggested that both isoforms populate highly disordered ensembles with nearly indistinguishable ensemble-averaged radii of gyration and end-to-end distances. In contrast, extensive all-atom simulations by Song *et al*. [31] reported that, while both peptides sample collapsed and extended states, A*β*42 exhibits an enhanced propensity to form compact, hydrophobically collapsed conformations with reduced hydrophobic SASA compared to A*β*40. Taken together, these apparently conflicting observations highlight that ensemble-averaged descriptors such as *R*_*g*_ and SASA are highly sensitive to force-field choices, modeling resolution, and analysis protocols, and may therefore mask substantial heterogeneity within the conformational ensemble. Rather than indicating an intrinsic ambiguity, this sensitivity underscores the need for analysis strategies capable of resolving recurrent conformational families beyond simple averages.

In this context, the clustering framework introduced in this study enables a more robust and physically interpretable comparison of the conformational landscapes of A*β*40 and A*β*42 by explicitly identifying and contrasting their dominant structural basins. The cluster-level analysis (Figures 6 and 6) confirms the trends inferred from global observables, showing that A*β*42 displays both a slightly higher degree of internal organization in *β*-rich states and a greater propensity to populate extended, solvent-exposed disordered basins compared to A*β*40.

Taken together, our results demonstrate that the addition of just two residues in A*β*42 is sufficient to bias the monomeric conformational ensemble toward a shifted balance between disorder and transient order. This shift is characterized by an increased propensity for *β*-like structures, a stronger involvement of the C-terminal region, and more frequent sampling of extended and solvent-exposed conformations. More broadly, our findings suggest that the divergent aggregation behaviors of A*β*40 and A*β*42 may originate from small but systematic redistributions of conformational populations already present at the monomeric level. Such subtle biases can be amplified during oligomerization and fibril formation, highlighting how minimal sequence variations in intrinsically disordered peptides can produce disproportionately large effects on collective behavior.

## Methods

### System Setup and Molecular Dynamics Simulations

All simulations of A*β*40 and A*β*42 were performed using identical protocols. The peptide sequences were taken from the Protein Data Bank (A*β*42: PDB ID 6SZF; A*β*40: PDB ID 1BA4). In both peptides, the C-terminal residue (A42 in A*β*42; V40 in A*β*40) was modeled with a deprotonated carboxylate group. The N-terminal aspartic acid carried a protonated amino group whose charge was neutralized by the negatively charged side chain, resulting in an overall neutral residue. Side-chain protonation states were assigned assuming physiological pH: lysine and arginine were kept protonated; non-terminal aspartic acid and glutamic acid were modeled in their deprotonated forms; and histidine was assigned as the neutral tautomer, which is the predominant state near physiological pH [53].

Each peptide was solvated in a cubic box of TIP3P water with an edge length of approximately 10 nm, under periodic boundary conditions in all directions. The box size was chosen to ensure at least 1.2–1.5 nm of solvent between the peptide and its periodic images, thereby excluding spurious inter-image interactions. All simulations employed the CHARMM36m force field [15], which provides an improved balance between secondary-structure propensities and intrinsic disorder relative to earlier CHARMM versions. CHARMM36m has been shown to reproduce conformational ensembles of intrinsically disordered peptides and proteins [15, 54, 32], and has been successfully used in previous studies of A*β* monomers and oligomers under physiological conditions [55].

All simulations were carried out using GROMACS [56] patched with PLUMED [57] for enhanced sampling. Production trajectories were generated in the NPT ensemble using a 2 fs time step. For each peptide, three independent replicas of 3 *µ*s were performed, yielding a total simulation time of 9 *µ*s per system. Coordinates were saved every 20 ps. The temperature was maintained at 310 K using the V-rescale thermostat [58], and pressure was maintained at 1 atm using the isotropic Parrinello–Rahman barostat [59, 60].

Electrostatics interactions were computed using Particle Mesh Ewald. Lennard-Jones interactions were treated using a force-switch scheme consistent with CHARMM36m: the van der Waals potential was smoothly switched between 1.0 and 1.2 nm, and the real-space cutoff for Coulomb interactions was set to 1.2 nm. Long-range dispersion corrections were disabled, as required for simulations using the CHARMM force field. Unless otherwise specified, simulations employed the default parameters of GROMACS (version 2024.3) and PLUMED (version 2.9.3).

### Collective Variables and Metadynamics Simulations

#### Collective variables

Enhanced sampling was performed using two collective variables (CVs) describing secondary structure content: the *α*- and *β*-content, implemented in PLUMED as ALPHARMSD and ANTIBETARMSD [38]. These CVs quantify the amount of *α*-helical and *antiparallel β*-sheet structure by comparing contiguous residue segments to ideal *α* and *β* blocks derived from a statistical analysis of protein structures in the PDB. Specifically:

- The *α*-content is obtained by evaluating all contiguous 6-residue segments and comparing them to the ideal *α*-helical block.
- The *β*-content is obtained by evaluating all contiguous 8-residue segments (with a two-residue separation between the central triplets) against the ideal antiparallel *β*-sheet block.

The similarity is quantified via the switching function

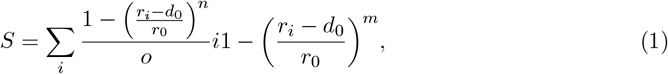

where *r*_*i*_ is the RMSD of each segment. Parameters were set to *d*_0_ = 0, *r*_0_ = 0.1, *n* = 8, and *m* = 12, as prescribed in Ref. [38].

Results are presented in terms of the CVs *n*_*α*_ and *n*_*β*_, obtained by rescaling the raw *α*- and *β*-content CVs by factors of 1.1 and 2.5, respectively. These rescaled quantities provide an approximate estimate of the number of residues engaged in *α*-helical or *β*-like structures, consistent with typical definitions used in secondary-structure assignment tools such as STRIDE [61] and DSSP [40]. The optimization of the rescaling factors and their validation against STRIDE and DSSP assignments are described in detail in the SI.

#### Metadynamics

Enhanced sampling was performed using well-tempered metadynamics (MetaD) [18]. Gaussian biases were deposited every 5 ps, with an initial height of 0.5 kJ*/*mol and an initial width of 0.1 (in CV units). The Gaussian width was chosen based on the natural fluctuations of the *α*- and *β*-content CVs, ensuring efficient filling of free-energy basins without oversmoothing the landscape. For computational efficiency, a grid in CV space was used to store the bias potential, with a resolution of 0.01. A bias factor of *γ* = 10 was selected to enhance sampling while restricting the exploration to conformations with physically reasonable free energies, thereby avoiding systematic visitation of highly unfavorable, non-representative states.

#### Free Energy Estimation

Free energy surfaces (FESs) were reconstructed from the MetaD bias using the well-tempered metadynamics (WT-MetaD) formalism [18], as implemented in the sum_hills utility of PLUMED [57]. In the well-tempered scheme, the time-dependent bias *V* (*ξ, t*) converges to a rescaled estimate of the underlying free energy:

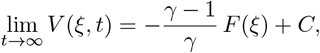

where *F* (*ξ*) is the physical free energy as a function of the collective variable *ξ* and *C* is an irrelevant additive constant. The bias is constructed as

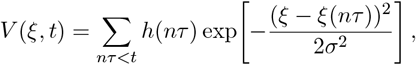

where *σ* is the Gaussian width and *h*(*nτ*) is the time-dependent height:

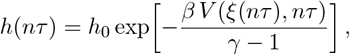

with *h*_0_ the initial Gaussian height and *β* = 1*/*(*k*_B_*T*).

Throughout the manuscript, FESs are expressed in terms of the structural variables *n*_*α*_ and *n*_*β*_ (see section *Collective Variables and Metadynamics Simulations*), which provide approximate estimates of the number of residues participating in *α*-helical or *β*-like structures.

#### Convergence assessment

Convergence was assessed by monitoring (i) the time evolution of the accumulated MetaD bias and (ii) the temporal stability of the relative populations associated with the dominant free energy minima. Convergence metrics reported in the Supporting Information show that bias deposition reaches a quasi-stationary regime and that the main FES features remain stable over the final portion of each trajectory, supporting the qualitative and comparative analyses presented for A*β*40 and A*β*42.

#### Reweighting Procedure

Because WT-MetaD changes the sampling distribution, unbiased ensemble averages can only be obtained through an explicit reweighting procedure. Here we adopt an estimator formally analogous to that used in umbrella sampling [62], adapted to the stationary distribution generated by WT-MetaD, for which the equilibrium probability density is proportional to exp[−*βF* (*ξ*)*/γ*] [18, 63].

To ensure that reweighting was applied only to configurations sampled under quasi-stationary bias conditions, the initial 1 *µ*s of each trajectory–during which the bias deposition had not yet stabilized–was excluded from all reweighted analyses. This choice provides improved numerical robustness compared with approaches based directly on the instantaneous bias, which can be particularly noisy for IDPs, which are characterized by broad and weakly funneled free-energy landscapes.

Unbiased observables were computed using the two-dimensional free-energy surface *F* (*α, β*) reconstructed from the combined sampling of the three independent replicas (i.e., the replica-averaged FES). Averaging across replicas mitigates statistical noise and reduces sensitivity to local convergence fluctuations in individual trajectories. For each configuration *i*, characterized by CV values (*α*_*i*_, *β*_*i*_) and an observable *x*_*i*_, the statistical weight was defined as

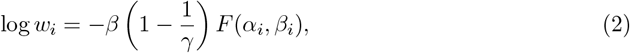

with *F* (*α*_*i*_, *β*_*i*_) obtained by bilinear interpolation over the replica-averaged FES grid. Because the FES spans tens of kJ*/*mol, we applied a standard numerical log-shift to prevent underflow:

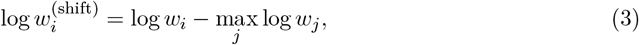

which preserves relative weights and does not affect ensemble averages. The reweighted histogram of an observable *x* was computed as

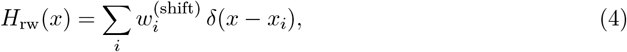

evaluated on a discrete grid. The final unbiased distribution was obtained by averaging per-replica histograms:

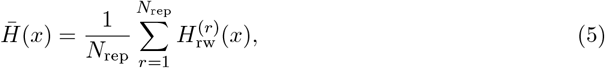

and statistical uncertainties are reported as the standard error of the mean across replicas.

This approach follows the theoretical framework of WT-MetaD [18] and its associated reweighting theory [63, 19], and provides numerically robust recovery of unbiased probability distributions from enhanced-sampling simulations.

### Secondary-structure analysis, solvent exposure, global compaction, and hydrogen bonding

Per-residue secondary-structure assignments were obtained using the DSSP algorithm [40], which classifies each residue at every frame into one of eight local conformations: 3_10_ helix, *α*-helix, *π*-helix, polyproline II, extended *β*-strand, isolated *β*-bridg, turn, and bend, with all remaining states reported as coil. For each frame, each `DSSP` label was encoded as a binary indicator and reweighted frame-by-frame as described in section *Reweighting Procedure*, yielding unbiased per-residue probabilities *p*_*i*_(*s*) for each DSSP state *s* at residue *i*.

Intra-peptide contact maps were computed using the MDAnalysis Python library [44]. Pairwise residue distances were calculated considering only C_*α*_ atoms, and a cutoff of 0.8 nm was applied to define residue–residue contacts. Contact probabilities were subsequently reweighted to obtain unbiased estimates.

Solvent-accessible surface area (SASA) was computed using `gmx sasa` [56], which implements the double-cubic lattice method [64]. Total SASA was decomposed into polar (P-SASA) and apolar (A-SASA) contributions following the Roseman hydrophobicity scale [48]: polar residues contribute to P-SASA, apolar residues to A-SASA, and glycine is excluded from both categories. The C-terminal residue, modeled as deprotonated in both A*β*40 and A*β*42, was treated as polar regardless of its Roseman classification. All SASA-based quantities were reweighted to recover unbiased ensemble distributions.

Global compaction was quantified via the radius of gyration *R*_*g*_, computed with `gmx polystat` [56] and subsequently reweighted.

Hydrogen bonds were evaluated using `gmx hbond` [56] with default angle and distance parameters, distinguishing intra-peptide (*n*_HB− IP_) and peptide–solvent hydrogen bonds (*n*_HB − PS_). Both time series were processed through the same reweighting procedure described above to obtain unbiased distributions.

### Conformational clustering

In STEP 1 of the clustering procedure (Figure 5), we used *n*_min_ = 10, *s*_min_ = 0.3, *N*_min_ = 300. *n*_*max*_ was set to 10 as for *n >* 10 we did not obtain acceptable clusters. The distance cutoff for local density calculation in DPC was initially set at 1.3 in the adimensional space of the four normalized descriptors, and then reduced by 10% at each iteration. The iteration on the density cutoff stops when either the remaining frames are less than *N*_min_, or no acceptable cluster had been generated in the last 5 iterations.

In STEP 2, clusters produced by different subsets and density scales were compared using the Wasserstein distances [65, 66]. In particular, as distances had to be calculated in the space of the four descriptors, we considered the modulus of the array composed by the four Wasserstein distances. The resulting distance matrix was subjected to agglomerative hierarchical clustering with average linkage [50]. The dendrogram was cut at the elbow of the merge-distance curve, yielding a unified set of *global clusters* consistent across all subsets.

#### Cluster relevance

As MetaD does not preserve unbiased state populations, raw cluster occupancies cannot be interpreted directly. To recover equilibrium populations, each frame was assigned a statistical weight *w*_*k*_ computed according to the reweighting formalism; see section *Reweighting Procedure*. The relevance of a global cluster *C*_*i*_ was defined as

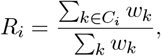

which represents the fraction of the unbiased equilibrium probability density associated with cluster *C*_*i*_. Accordingly, a cluster with high *R*_*i*_ corresponds to a structurally coherent and frequently sampled basin in the unbiased ensemble, whereas clusters with low *R*_*i*_ reflect rare or transient conformations.

Overall, this framework integrates density-based detection, hierarchical merging, and replica-aware reweighting, yielding a statistically robust and physically interpretable partition of the A*β*40 and A*β*42 conformational landscapes.

## Notes

### Competing Interest Statement

The authors have declared no competing interest.

### Funding

- Ministero dell’università e della ricerca: PRIN 2022 - Code: 20222YMAR5 - Project title: NeuRaftAmyl

