## Supporting Information for "The monomeric conformational ensembles of A*β*40 and A*β*42 encode their differential amyloid aggregation propensity"

### Supporting Information
### The monomeric conformational ensembles of A$\beta$40 and A$\beta$42 encode their differential amyloid aggregation propensity

Irene Cadenelli[1], Andrea Ciccolo[2], Andrea Tagliabue[1], Giulia Rossi[1], Valeria Conti-Nibali[2,*], and Davide Bochicchio[1,*]

[1] *Department of Physics, University of Genoa, Genoa, Italy*
[2] *Department of Mathematical and Computational Sciences, Physical Sciences and Earth Sciences, University of Messina, Messina, Italy*

#### 1 Supporting

##### 1.1 Convergence of Well-Tempered Metadynamics Simulations

Convergence of the WT-MetaD simulations [1] was assessed by monitoring the time evolution of a free energy difference between two selected regions of the one-dimensional FES projections along the biased collective variables $n_\alpha$ and $n_\beta$. The reported curves correspond to replica-averaged quantities obtained by combining the three independent simulations performed for each peptide (Fig. 1). For each CV, two windows were defined in the rescaled variables: a low-$n$ region (0–5 residues), enriched in compact, weakly structured conformations, and a high-$n$ region (5–30 residues), associated with more extended and highly structured states. The resulting $\Delta FE(t)$ provides a practical and sensitive indicator of sampling stability in enhanced-sampling simulations of intrinsically disordered peptides and amyloid-$\beta$ monomers [1, 2].

In a converging WT-MetaD simulation, $\Delta FE(t)$ is expected to display damped oscillations and to approach a quasi-stationary regime as the accumulated bias becomes effectively time-independent. As shown in Fig. 1, both A$\beta$40 and A$\beta$42 exhibit decreasing oscillation amplitudes and fluctuations around slowly drifting mean values for both $n_\alpha$ and $n_\beta$. Residual long-time fluctuations are intrinsic to disordered systems, whose rugged and weakly funneled landscapes hinder strict numerical convergence on accessible simulation timescales, particularly for amyloid-$\beta$ monomers [3, 4].

To ensure robust equilibrium estimates from reweighting, the initial 1 $\mu$s of each replica was excluded from all reweighted analyses, thereby removing the early transient regime characterized by rapid bias deposition. Reweighting was performed using standard WT-MetaD formalisms that reconstruct the equilibrium Boltzmann distribution from the biased dynamics [5]. After this equilibration interval, the overall FES topology and the relative stability of the low- and high-content *alpha* add $\beta$ regions remain stable over extended time windows, supporting the reliability of the reweighted equilibrium ensemble.

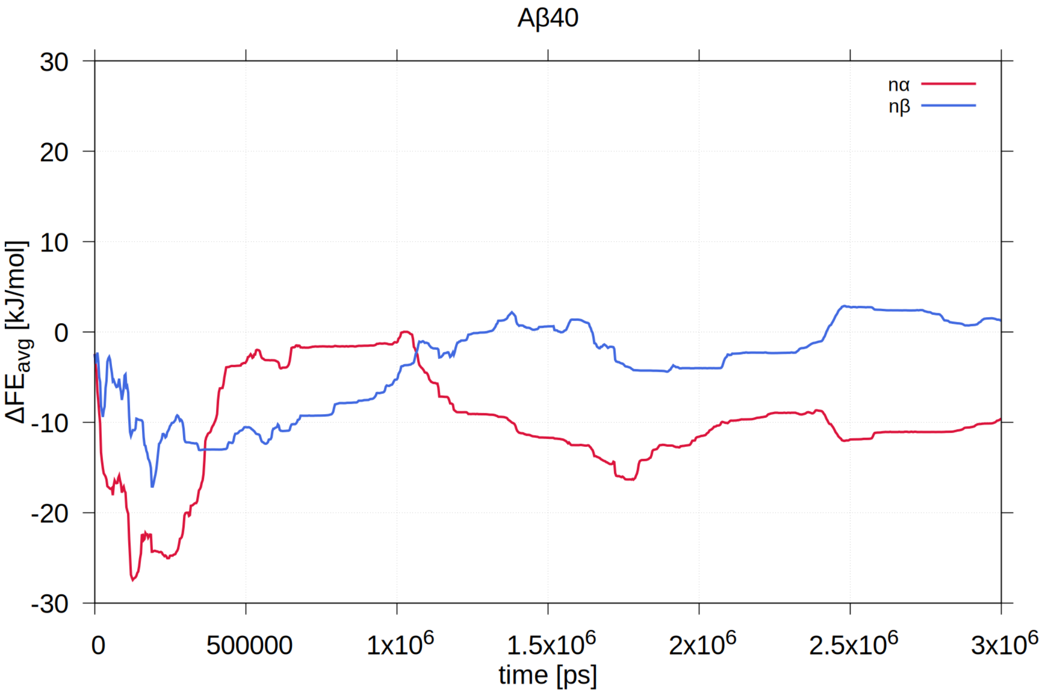


**(a)** A$\beta$40

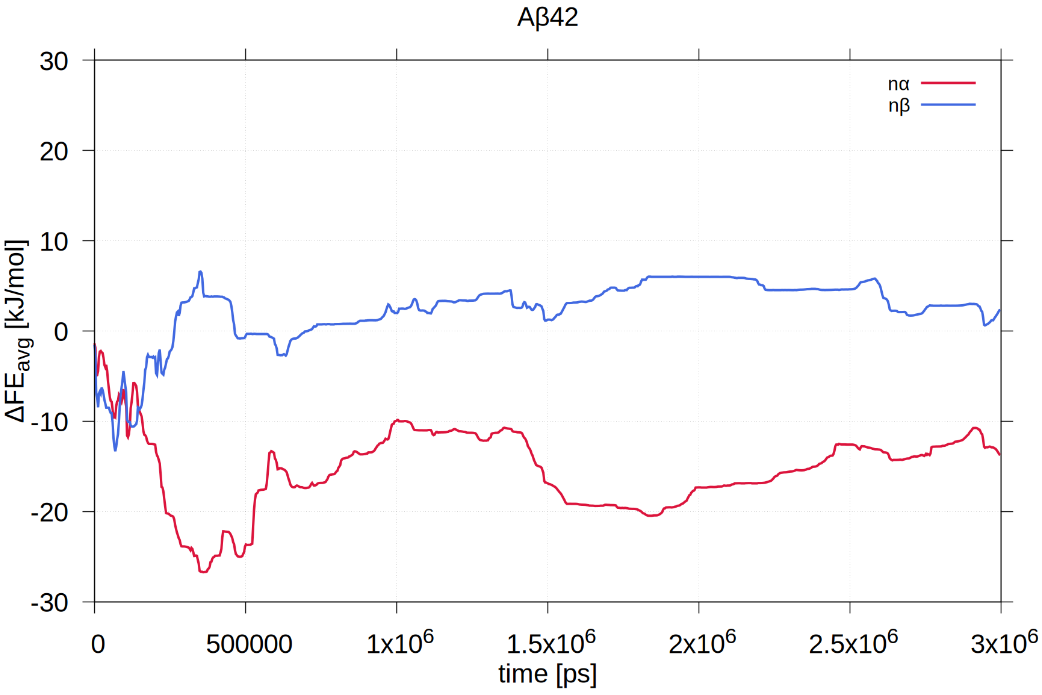


**(b)** A$\beta$42

**Figure 1: Convergence of replica-averaged WT-MetaD simulations.** Time evolution of the replica-averaged free energy difference $\Delta FE_{\mathrm{avg}}$ between low-$n$ (0–5 residues) and high-$n$ (5–30 residues) regions of the one-dimensional FES projections along $n_\alpha$ (red) and $n_\beta$ (blue) for (a) A$\beta$40 and (b) A$\beta$42. Free-energy differences were estimated from WT-MetaD using standard estimators for converged bias potentials [1, 2].

#### 1.2 Validation of the PLUMED-Based Secondary Structure Estimators

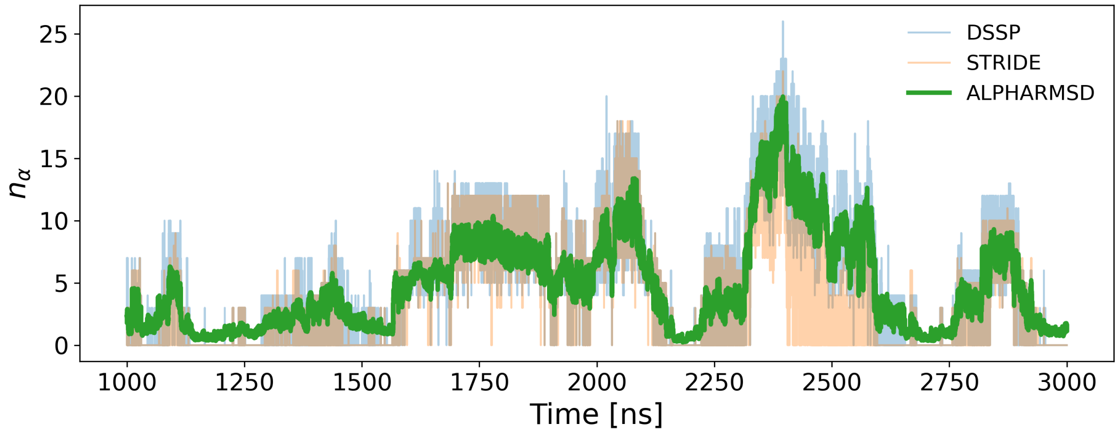


**(a)** Comparison of $n_\alpha$ estimates for A$\beta$42 (replica 1).

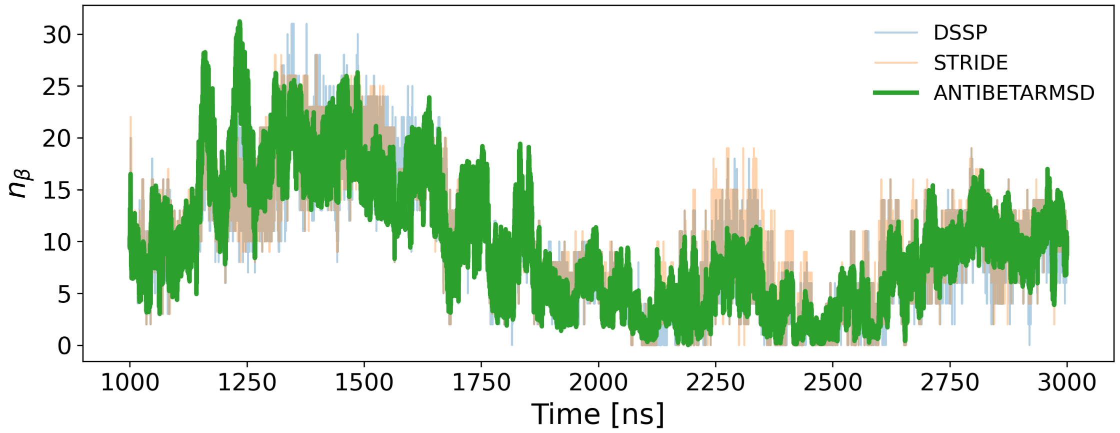


**(b)** Comparison of $n_\beta$ estimates for A$\beta$42 (replica 1).

**Figure 2: Validation of rescaled PLUMED collective variables for A$\beta$42.** Time-series comparison between secondary structure content estimated using DSSP and STRIDE and the corresponding PLUMED CVs after rescaling. Panel (a) reports the number of residues in $\alpha$-helical structure from DSSP, STRIDE, and the rescaled PLUMED variable ALPHARMSD ($n_\alpha = c_\alpha \cdot$ ALPHARMSD). Panel (b) reports the analogous comparison for $\beta$-strand content using ANTIBETARMSD ($n_\beta = c_\beta \cdot$ ANTIBETARMSD).

The PLUMED CVs `ALPHARMSD` and `ANTIBETARMSD` [6] provide continuous measures of $\alpha$-helical and $\beta$-strand content, respectively, but are not directly interpretable as residue counts. To obtain physically meaningful residue-level estimates, we used established secondary structure assignment methods, namely DSSP [7] and STRIDE [8], which are routinely adopted as reference estimators for amyloid-$\beta$ peptides and other IDPs [9, 4].

To establish a quantitative correspondence between PLUMED-derived observables and residue counts, multiplicative rescaling factors $c_\alpha$ and $c_\beta$ were calibrated against DSSP and STRIDE assignments for A$\beta$42. For each of the three independent replicas, PLUMED-derived $n_\alpha$ and $n_\beta$ were compared with the corresponding DSSP and STRIDE values over the 1.0–3.0 $\mu$s time interval, and optimal rescaling factors were determined by minimizing the least-squares deviation between the methods.

The resulting rescaling factors showed excellent consistency across replicas. Averaging over the three estimates yielded $c_\alpha = 1.1$ and $c_\beta = 2.5$. These values were subsequently used throughout the study to convert PLUMED outputs into estimates of the number of residues participating in $\alpha$-helical and $\beta$-strand structure. Representative time-series comparisons for replica 1 are shown in Fig. 2, demonstrating close agreement between the rescaled PLUMED variables and the DSSP and STRIDE references.

##### 1.3 Complete Cluster Characterization

Figures S3–6 provide a comprehensive cluster-resolved characterization of the conformational ensembles of A$\beta$40 and A$\beta$42 obtained from the consensus clustering analysis. Such a detailed description is particularly relevant for amyloid-$\beta$ peptides, whose monomeric ensembles are intrinsically heterogeneous and span a wide range of structural and physicochemical states, even at equilibrium conditions [3, 4]. For each identified cluster, the reported tables include:

- Reweighted equilibrium population (%).
- Primary clustering descriptors ($n_\alpha$, $n_\beta$, $R_g$, total SASA): weighted mean ± weighted standard deviation.
- Post hoc physicochemical descriptors (A-SASA, P-SASA, $n_{\text{HB-IP}}$, $n_{\text{HB-PS}}$): weighted mean ± weighted standard deviation.
- Global reference values (`ALL`): corresponding reweighted averages over the full unbiased ensemble.

Together, these data provide a complete numerical summary of all conformational families identified in the analysis. The cluster-resolved descriptors complement the qualitative discussion of the dominant conformational states reported in the main text and enable a direct quantitative comparison between A$\beta$40 and A$\beta$42 at the level of individual free-energy basins.

| Rank | Relevance [%] | nα | nβ | Rg [nm] | SASA [nm²] |
|---|---|---|---|---|---|
| 1 | 55.4 | 2.55 ± 1.48 | 9.65 ± 1.96 | 1.08 ± 0.0599 | 36.7 ± 1.89 |
| 2 | 11.8 | 1.95 ± 1.5 | 11.8 ± 2.4 | 1.3 ± 0.141 | 41.5 ± 1.67 |
| 3 | 8.3 | 3.38 ± 1.66 | 2.49 ± 1.63 | 1.3 ± 0.116 | 44.1 ± 1.81 |
| 4 | 7.09 | 1.86 ± 1.21 | 2.37 ± 1.82 | 1.61 ± 0.149 | 49.1 ± 2.14 |
| 5 | 6.92 | 4.18 ± 1.71 | 4.55 ± 1.72 | 1.14 ± 0.0918 | 39.2 ± 1.82 |
| 6 | 4.37 | 0.776 ± 0.588 | 9.45 ± 2.17 | 1.77 ± 0.216 | 47.2 ± 2.11 |
| 7 | 3.98 | 1.56 ± 1.7 | 1.75 ± 2.15 | 2.2 ± 0.273 | 53.1 ± 2.18 |
| 8 | 0.975 | 2.41 ± 1.45 | 18.2 ± 3.62 | 1.1 ± 0.101 | 35.2 ± 1.98 |
| 9 | 0.796 | 9.19 ± 1.51 | 2.53 ± 1.51 | 1.2 ± 0.0969 | 40.2 ± 2.07 |
| 10 | 0.222 | 8.54 ± 2.2 | 0.849 ± 0.91 | 1.59 ± 0.185 | 47 ± 2.12 |
| 11 | 0.0768 | 8.49 ± 1.47 | 7.67 ± 2.15 | 1.08 ± 0.0816 | 36 ± 1.71 |
| 12 | 0.0213 | 13.9 ± 1.66 | 2.83 ± 1.61 | 1.07 ± 0.0838 | 36.4 ± 1.77 |
| 13 | 0.00373 | 15.5 ± 1.43 | 1.76 ± 1.47 | 1.25 ± 0.126 | 39.8 ± 1.89 |
| 14 | 0.000469 | 12.9 ± 1.6 | 6.83 ± 1.88 | 0.984 ± 0.0414 | 33.2 ± 1.32 |
| 15 | 0.000405 | 9.13 ± 1.9 | 12.7 ± 2.29 | 1.34 ± 0.138 | 39.9 ± 1.65 |
| 16 | 0.000227 | 17.9 ± 2.84 | 0.822 ± 0.607 | 1.5 ± 0.137 | 43.8 ± 1.86 |
| 17 | 0.000166 | 19.7 ± 2 | 1.48 ± 0.963 | 1.12 ± 0.135 | 36.2 ± 2.17 |
| ALL | | 6.27 ± 5.3 | 7.39 ± 6.02 | 1.21 ± 0.256 | 39 ± 4.82 |

**Figure 3: Cluster-resolved structural descriptors for A$\beta$40.** Reweighted equilibrium populations and weighted averages of $n_\alpha$, $n_\beta$, radius of gyration $R_g$, and total SASA for all consensus clusters.

| Rank | Relevance [%] | nα | nβ | Rg [nm] | SASA [nm²] |
|---|---|---|---|---|---|
| 1 | 56.2 | 2.74 ± 1.75 | 12 ± 2.17 | 1.14 ± 0.0749 | 38.1 ± 2.03 |
| 2 | 11.3 | 2.89 ± 1.64 | 2.85 ± 1.85 | 1.42 ± 0.142 | 46 ± 2.25 |
| 3 | 9.71 | 0.936 ± 0.602 | 7.58 ± 2.11 | 1.57 ± 0.205 | 47.5 ± 2.3 |
| 4 | 8.12 | 3.87 ± 1.57 | 5.17 ± 2.24 | 1.17 ± 0.0945 | 40.7 ± 2.11 |
| 5 | 6.95 | 1.25 ± 1.08 | 1.44 ± 1.77 | 2.03 ± 0.307 | 54.6 ± 2.6 |
| 6 | 4.46 | 4.2 ± 1.91 | 10.8 ± 2.47 | 1.47 ± 0.126 | 43.6 ± 1.4 |
| 7 | 2.47 | 1.61 ± 1.33 | 16.5 ± 2.39 | 1.51 ± 0.171 | 42.7 ± 1.61 |
| 8 | 0.499 | 2.36 ± 1.55 | 20.5 ± 3.18 | 1.18 ± 0.0868 | 36.7 ± 1.66 |
| 9 | 0.194 | 7.14 ± 1.52 | 6.49 ± 3.06 | 1.06 ± 0.0619 | 36.5 ± 2.19 |
| 10 | 0.0699 | 5.84 ± 2.51 | 0.446 ± 0.888 | 2.29 ± 0.338 | 54.6 ± 2.67 |
| 11 | 0.00967 | 10.7 ± 1.9 | 3.02 ± 2.18 | 1.14 ± 0.109 | 39.3 ± 3.22 |
| 12 | 0.000699 | 11.4 ± 2.15 | 1.2 ± 1.44 | 1.46 ± 0.154 | 46.9 ± 2.15 |
| 13 | 5.32e-08 | 15.3 ± 2.14 | 4.05 ± 2.6 | 1.38 ± 0.14 | 42.7 ± 2.22 |
| ALL | | 4.6 ± 3.42 | 8.59 ± 6.56 | 1.26 ± 0.277 | 40.6 ± 5.18 |

**Figure 4: Cluster-resolved structural descriptors for A$\beta$42.** Same quantities as in Fig. 3, reported for the A$\beta$42 conformational ensemble.

| Rank | Relevance [%] | A-SASA [nm²] | P-SASA [nm²] | nHB-IP | nHB-PS |
|---|---|---|---|---|---|
| 1 | 55.4 | 14 ± 1.74 | 20.5 ± 2.02 | 15.5 ± 3.35 | 123 ± 8.14 |
| 2 | 11.8 | 16.5 ± 1.85 | 22.2 ± 1.81 | 14.3 ± 2.74 | 128 ± 7.03 |
| 3 | 8.3 | 18 ± 1.72 | 23 ± 1.72 | 10 ± 3.01 | 136 ± 7.68 |
| 4 | 7.09 | 20.4 ± 1.84 | 25.2 ± 1.72 | 7.45 ± 2.8 | 143 ± 7.63 |
| 5 | 6.92 | 15.4 ± 1.9 | 21.3 ± 1.92 | 12.8 ± 3.01 | 128 ± 7.47 |
| 6 | 4.37 | 19.6 ± 1.61 | 24.4 ± 1.92 | 10.1 ± 2.81 | 139 ± 7.66 |
| 7 | 3.98 | 22.6 ± 1.71 | 27 ± 1.75 | 5.72 ± 2.41 | 148 ± 7.4 |
| 8 | 0.975 | 13.1 ± 1.49 | 19.9 ± 1.49 | 18.9 ± 3.15 | 116 ± 7.34 |
| 9 | 0.796 | 15.6 ± 1.77 | 22.1 ± 1.72 | 14.6 ± 3.17 | 125 ± 7.45 |
| 10 | 0.222 | 19.1 ± 1.59 | 24.6 ± 1.6 | 11.6 ± 2.97 | 133 ± 7.27 |
| 11 | 0.0768 | 13.9 ± 1.57 | 20.1 ± 1.69 | 17.9 ± 3.2 | 117 ± 7.3 |
| 12 | 0.0213 | 13.4 ± 1.89 | 20.5 ± 1.59 | 18.6 ± 3.4 | 116 ± 7.18 |
| 13 | 0.00373 | 15.1 ± 1.16 | 22.2 ± 1.35 | 17.9 ± 3.28 | 119 ± 7.52 |
| 14 | 0.000469 | 12.4 ± 1.77 | 18.4 ± 1.47 | 21.1 ± 3.32 | 110 ± 6.83 |
| 15 | 0.000405 | 14.9 ± 1.32 | 22.8 ± 1.26 | 18.5 ± 2.98 | 119 ± 6.53 |
| 16 | 0.000227 | 17.4 ± 1.46 | 23.6 ± 1.63 | 17.9 ± 3.63 | 121 ± 8.14 |
| 17 | 0.000166 | 13.7 ± 1.26 | 20.3 ± 1.44 | 21.3 ± 3.08 | 110 ± 6.87 |
| ALL | | 15.2 ± 2.88 | 21.4 ± 2.46 | 15.4 ± 4.96 | 124 ± 11.8 |

**Figure 5: Extended physicochemical descriptors for A$\beta$40.** Cluster-resolved apolar and polar SASA and hydrogen-bond metrics (nHB-IP and nHB-PS).

| Rank | Relevance [%] | A-SASA [nm²] | P-SASA [nm²] | nHB-IP | nHB-PS |
|---|---|---|---|---|---|
| 1 | 56.2 | 14.1 ± 1.67 | 21.5 ± 1.31 | 16.4 ± 3.29 | 125 ± 8.04 |
| 2 | 11.3 | 18.9 ± 1.65 | 24.2 ± 1.52 | 10 ± 2.83 | 140 ± 7.55 |
| 3 | 9.71 | 19.3 ± 1.49 | 25.2 ± 1.75 | 10.3 ± 2.72 | 141 ± 7.62 |
| 4 | 8.12 | 15.7 ± 1.84 | 22.3 ± 1.48 | 13.5 ± 2.89 | 131 ± 7.38 |
| 5 | 6.95 | 23.6 ± 2.1 | 27.3 ± 1.36 | 5.58 ± 2.48 | 152 ± 7.61 |
| 6 | 4.46 | 18 ± 1.13 | 23.2 ± 1.21 | 14.3 ± 2.96 | 132 ± 7.18 |
| 7 | 2.47 | 17.2 ± 0.971 | 22.9 ± 1.41 | 15.5 ± 2.74 | 131 ± 7.2 |
| 8 | 0.499 | 14 ± 1.79 | 20.2 ± 1.49 | 20.3 ± 3.05 | 118 ± 7.66 |
| 9 | 0.194 | 13.1 ± 1.73 | 20.9 ± 1.32 | 18 ± 3.33 | 120 ± 7.67 |
| 10 | 0.0699 | 25 ± 1.38 | 26 ± 1.5 | 7.84 ± 3.31 | 148 ± 8.54 |
| 11 | 0.00967 | 15.8 ± 2.23 | 21.1 ± 1.78 | 17 ± 4 | 123 ± 9.19 |
| 12 | 0.000699 | 20.3 ± 1.67 | 23.7 ± 1.3 | 13.6 ± 3.07 | 133 ± 7.07 |
| 13 | 5.32e-08 | 17.8 ± 1.38 | 22.6 ± 1.35 | 19.2 ± 2.83 | 123 ± 6.81 |
| ALL | | 15.9 ± 3.28 | 22.1 ± 2.26 | 15.4 ± 4.89 | 128 ± 11.9 |

**Figure 6: Extended physicochemical descriptors for A$\beta$42.** Same quantities as in Fig. 5, reported for the A$\beta$42 conformational ensemble.
